# Genome-wide identification and comparative analysis of *Dof* gene family in *Brassica napus*

**DOI:** 10.1101/2020.12.15.422814

**Authors:** Neeta Lohani, Saeid Babaei, Mohan B. Singh, Prem L. Bhalla

**Affiliations:** Plant Molecular Biology and Biotechnology Laboratory, Faculty of Veterinary and Agricultural Sciences, The University of Melbourne, Parkville, Melbourne, VIC 3010, Australia

**Author notes:** Corresponding author: Prem L Bhalla.

**Keywords:** *Dof*, *Brassica napus*, canola, Transcription factor, Polyploidy, Abiotic stress

## Abstract

DOF, DNA binding with one finger proteins are plant-specific transcription factors shown to play roles in diverse plant functions. However, a—little is known about DOF protein repertoire of the allopolyploid crop, *Brassica napus*. Here, we report genome-wide identification and systematic analysis of the *Dof* transcription factor family in this important oilseed crop. We identified 117 *Brassica napus* Dof genes (*BnaDofs*). So far, this is the largest number of *Dof* genes reported in a single eudicot species. Based on phylogenetic analysis, *BnaDofs* were classified into nine groups (A, B1, B2. C1, C2.1, C2.2, C3, D1, D2). Most members belonging to a particular group displayed conserved gene structural organisation and similar protein motifs distribution. Chromosomal localisation analysis highlighted the uneven distribution of *BnaDofs* across all chromosomes. Evolutionary analysis exemplified that the divergence of *Brassica* genus from Arabidopsis, the whole genome triplication event, and the hybridisation of *B. oleracea* and *B. rapa* to form *B. napus*, followed by gene loss and rearrangements, led to the expansion and divergence of Dof TF gene family in *B. napus*. Functional annotation of BnaDof proteins, cis-element analysis of their promoters suggested potential roles in organ development, the transition from vegetative to the reproductive stage, light responsiveness, phytohormone responsiveness as well as abiotic stress responses. Furthermore, the transcriptomic analysis highlighted the preferential tissue-specific expression patters of *BnaDofs* and their role in response to various abiotic stress. Overall, this study provides a comprehensive understanding of the molecular structure, evolution, and potential functional roles of *Dof* genes in plant development and abiotic stress response.

## Introduction

*Brassica napus*, the second largest economically important oilseed crop is used as edible oil, livestock forage and in pharmaceuticals, cosmetics, and biofuel industry (USDA 2020). The yield of *B. napus* is constrained by harsh environmental conditions such as drought, extreme temperature, and salinity (Lohani et al. 2020). Dissecting the evolution and function of diverged plant specific transcription factor (TF) families such as DOF (DNA binding with one finger) is required for gaining the fundamental knowledge about the mechanisms underlying stress responses in *B. napus* and for developing stress-tolerant varieties for climate-smart agriculture.

The DOF TFs are plant specific transcription factors, first identified in maize in 1995 and was shown to play an essential role in regulating carbon metabolism related and light-regulated genes (Yanagisawa 1995; Yanagisawa 2000; Yanagisawa and Sheen 1998). Since then, diverse number of Dof TFs have been identified in several plants including 36 in Arabidopsis, 30 in rice up to 96 in wheat (Lijavetzky et al. 2003; Liu et al. 2020; Yanagisawa 2002). The Arabidopsis studies showed the first protein-protein interaction between a Dof domain protein with bZIP proteins associated with stress responses indicating this TF function in complex regulatory networks (Tamai et al. 2002).

Dof TFs are typically composed of 200-400 amino acid residues with a variable C-terminal region and, mainly characterized by a highly conserved DNA-binding domain i.e., the Dof domain, located towards the N-terminal of the protein (Le Hir and Bellini 2013; Yanagisawa 2004). Dof domain consists of only one Cys2/Cys2 zinc finger structure, which includes about 52 amino acid residues that specifically recognizes and binds a cis-regulatory element (5’-T/AAAAG-3’) in the gene promoters (Umemura et al. 2004) Another specific feature of Dof TFs is a bipartite nuclear localization signal (NLS) which is comprised of two short basic flanking regions (B1 and B2) with a 17 amino acid spacer. The bipartite NLS is highly conserved and plays a vital role in directing Dof TFs to the cell nucleus (Krebs et al. 2010).

Phylogenetic studies suggest a common ancestor (conserved as a single copy in *Chlamydomonas*) of *Dof* genes, which through numerous rounds of gene duplication drove the structural and thus functional diversification of the *Dof* gene family (Umemura et al. 2004). This evolutionary diversification may be related to acquiring new specialised functions needed to adjust and adapt to diverse and complex plant growth conditions.

*Brassica napus*, an amphidiploid (AACC, 2n=4x=38), originated from natural crossing between two ancestral diploid parents, *B. oleracea* (CC, 2n=2x=18) and *B. rapa* (AA, 2n=2x=20), about 7,500 years ago (Chalhoub et al. 2014). *Brassica* species belonging to the *Brassica* genus and Brassicaceae family offer a valuable model for studying polyploid genome evolution, mechanisms associated with gene duplications, loss of duplicated genes and gene neo- and sub-functionalization (Cheng et al. 2018; Liu et al. 2014). The availability of *B. napus, B. rapa* and *B. oleracea* genome sequences has provided an exceptional opportunity to identify and characterize key genes from a genome-wide perspective (Chalhoub et al. 2014; Liu et al. 2014; Wang et al. 2011).

This study aims to provide a complete description of the *Dof* gene family in *B. napus* by performing a genome-wide *in-silico* identification, characterisation, evolutionary and functional analysis of the *Dof* gene family. We report identification of one hundred seventeen genes as members of the *Dof* transcription factor family from *B. napus* belonging to nine groups. A detailed analysis of the *Dof* genes in terms of physical properties of proteins, chromosomal location, gene structure, motif analysis and phylogenetic relationships, and gene duplication was also performed. Orthology and synteny analysis were also carried out to explore the evolutionary history and divergence of the *Dof* TFs family in *B. napus, B. rapa, B. oleracea* and Arabidopsis. Further, functional annotation, cis-acting regulatory element analysis of the promoters of *BnaDof* genes along with their expression profiles highlighted their potential roles in regulating distinct and diverse developmental processes and stress responses.

## Materials and Methods

### Identification of *Dof* gene family members in *B. napus*

BLASTP search of the *B. napus* proteome was carried out using zf-Dof-domain search model accession (Pfam: PF02701) as a query to obtain the consensus amino-acid sequences of the putative Dof proteins. The term ‘Dof’ and Dof-domain search model accession ‘PF02701’ were used to search the Plant Transcription Factor Database 4.0 database (PlantTFDB) (Jin et al. 2016). To identify integrated Dof domain in the putative Dofs obtained from BLASTP and PlantTFDb search; SMART 8.0 software (http://smart.embl-heidelberg.de/) was used and the final predicted Dofs were further characterised (Letunic and Bork 2018). Expasy server’s ProtParam tool (https://web.expasy.org/protparam/) was used to compute the various physical and chemical properties of the predicted Dof proteins such as number of amino acids in the protein sequence, and molecular weight (Mw), protein isoelectric point (pI) and GRAVY (Grand Average of Hydropathy) of the protein (Gasteiger et al. 2005). Chromosomal locations as well as the genomic, coding, peptide, and promoter sequences of the *Dof* TFs were downloaded from GenoScope (Brassica napus.annotation_v5) database. Unique gene identifiers were assigned to the *Dofs* and they were referred to as BnaDofs.

### Evolutionary and gene duplication analysis of BnaDofs

Multiple Sequence alignments were performed on the Dof amino acid sequences using CLUSTALW with default settings (Larkin et al. 2007). MEGA7.0 was used to construct a phylogenetic tree (unrooted) based on the neighbour-joining method and pairwise gap deletion parameters. Statistical support for each tree node was provided by performing 1000 replicates bootstrap analysis (Kumar et al. 2008). Poisson correction method was employed to compute the evolutionary distances.

To identify gene duplications in *BnaDofs*, all *B. napus* gene sequences (101040) were first aligned using BLASTp, with an e-value of 1e-10 and then the duplication patterns were classified into interspersed and tandem duplications with MCScanX (default parameters) (Korf et al. 2003; Wang et al. 2012). Gene duplication was also analysed based on sequence similarity criteria i.e., similarity of the aligned regions of protein ≥ 80% (Lohani et al. 2019). Evolutionary analyses were conducted in MEGA7 (Kumar et al. 2008). The number of synonymous substitutions per synonymous site (dS/Ks), and the number of non-synonymous substitutions per non-synonymous site (dN/Ka), were calculated using the Nei-Gojobori method (Jukes-Cantor. The formula T□=□Ks/2R (where, Ks= number of synonymous substitutions per synonymous site, R=1.5□×□10^-8^ synonymous substitutions per site per year, T= divergence time) was used to estimate divergence time (Koch et al. 2000; Nei and Gojobori 1986).

### Gene structure and motif analysis of *BnaDofs*

The gene structure in terms of exon-intron organisation was determined using the GSDS2.0 (Gene Structure Display Server; http://gsds.cbi.pku.edu.cn) (Hu et al. 2015). MEME tool from the MEME suite 5.1.1 (http://meme-suite.org/tools/meme) was used to identify fifteen statistically significant motifs of the BnaDof protein sequences based on “zero or one occurrence per sequence (zoops)” (Bailey et al. 2015). The discoverable motif length and sites were set to 6-50 and 2–600, respectively.

### Functional annotation and promoter analysis

Functionally annotation of BnaDofs was performed using PANNZER2 (Protein ANNotation with Z-scoRE 2), which provides both Gene Ontology (GO) annotations and free text description predictions (Törönen et al. 2018). Promoter analysis of the *BnaDof* genes was performed by using the PlantCARE database (http://bioinformatics.psb.ugent.be/webtools/plantcare/html/and peptide sequences were downloaded from) to identify the cis-acting regulatory elements with putative involvement in various abiotic stress responses (Lescot et al. 2002). 1500□bp upstream regions from the start codon (ATG) of the *Dof* genes were downloaded as the promoter sequences from the EnsemblPlants Database (Bolser et al. 2016).

### Orthology and collinearity analysis of BnaDofs

Genome annotations and peptide sequences were downloaded from GenoScope (Brassica_napus.annotation_v5) and EnsemblPlants (Brassica_rapa.IVFCAASv1.36, Brassica_oleracea.v2.1.36). The orthologous genes in *B. oleracea, B. rapa, B. napus* and *A. thaliana* were identified using OrthoVenn2 (https://orthovenn2.bioinfotoolkits.net/home) (Xu et al. 2019).For the analysis of collinearity/synteny, 58 out of 117 Dof transcription factors members identified in *B. napus* where discarded because their chromosome of origin is uncertain. Syntenic blocks have been identified by sequence similarity search of the remaining 59 Dof members in B. napus against the reference genomes of Arabidopsis, *B. rapa* and *B. oleracea*, using blastn v. 2.10.1+ with stringent parameters (cutoff e-value 10e-50 and minimum percentage of identity of 75) (Korf et al. 2003). A custom python script was used to reformat the blast output and identify collinear genes and genes that underwent chromosomal translocation. Synteny plots were plotted with circos v. 0.69-9 (Krzywinski et al. 2009).

### Tissue-Specific expression and abiotic stress response expression profiling of BnaDofs

To investigate tissue specific expression (under non-stressed conditions) and abiotic stress responsive expression of the *BnaDof* genes, RNASeq data sets from previously published literature were downloaded from NCBI Sequence Read Archive database. Transcript expression was quantified using Kallisto v0.44.0 and read abundance was expressed as Transcripts Per Kilobase Million (TPM) (Bray et al. 2016). Heat maps were drawn by using ComplexHeatmap package to visualise the expression of *BnaDofs* (Gu 2015). Additionally, the expression maps for *BnaDofs* were downloaded from Brassica expression Data Base (BrassicaEDB). The Supplementary Folder with the downloaded expression maps of *BnaDofs* is accessible via the following link https://jmp.sh/odBHlfT.

## Results

### Identification and characterisation of *Brassica napus* DOF gene family

To identify the Dof TF family genes in *B. napus*, we carried out BLASTP searches using Dof-domain search model accession (Pfam: PF02701), searched the PlantTFDB4.0 and finally retrieved and analysed 156 amino acid sequences using the SMART8.0 database (Jin et al. 2016; Korf et al. 2003; Letunic and Bork 2018). We identified 117 full-length *B. napus* genes as putative members of the *Dof* gene family from *Brassica napus* reference genome (Brassica_napus.annotation_v5). We identified 117 full-length *B. napus* genes as putative members of the *Dof* gene family from *Brassica napus* reference genome *(Brassica_napus*. annotation_v5). We assigned new identifiers to the 117 *B. napus Dof* genes by using the prefix “*Bna*” for *B. napus* followed by *“Dof”* and a number based on their chromosomal locations (Figure 1). Thus, the identified *B. napus Dof* family members were named as *BnaDof01* to *BnaDof117* (Table S1). The *BnaDofs* were distributed on all 19 chromosomes, with 60 *BnaDofs* located on A genome (48 on chromosomes A01-A10), 56 located on C genome (40 on chromosomes C01-C09) and one gene placed on an unknown chromosome (*BnaDof117; BnaUnng03510D*). Chromosome C03 being the largest included the most *BnaDofs* (11 *BnaDofs*); followed by A09 with 10 *BnaDofs*. Out of 117 *BnaDof* genes, the exact chromosomal location of 58 genes was unknown.

**Figure 1.**
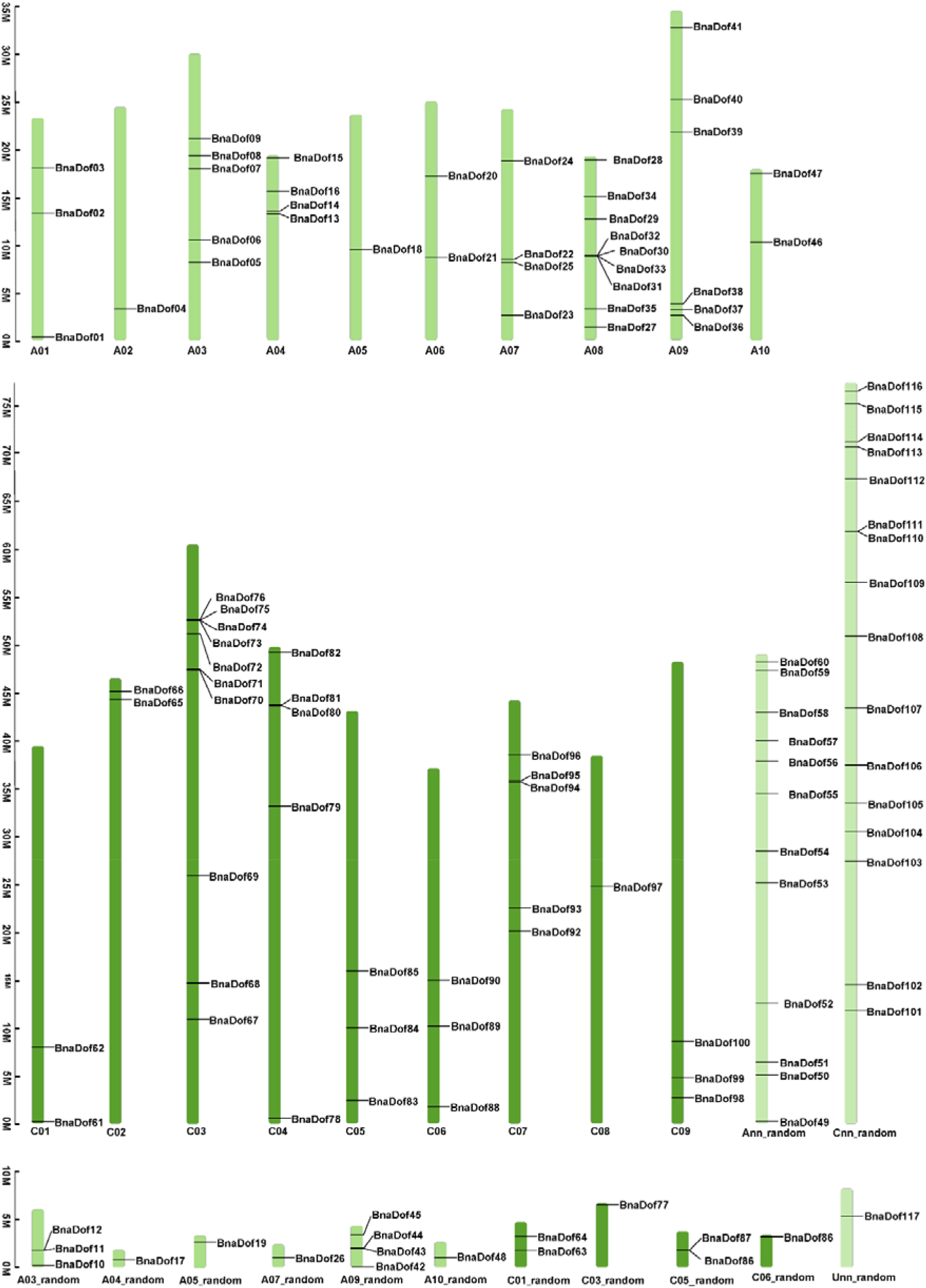
Chromosomal mapping and frequency distribution of *Brassica napus* DOF transcription factors (*BnaDofs01-117*).

The physical and chemical properties of BnaDof proteins are outlined in Table S1. The size of the BnaDof protein sequences ranged from 77 (BnaDof30, BnaDof31, BnaDof75, BnaDof76) to 453 (BnaDof33) amino acids. All BnaDofs had a marginally higher percentage of aliphatic amino acid than aromatic amino acids. The predicted isoelectric point (pI) ranged from 4.78 (BnaDof20) to 10.41 (BnaDof30, BnaDof31, BnaDof75, BnaDof76), and the molecular weight (MW) ranged from 8.76□kD (BnaDof30, BnaDof31, BnaDof75, BnaDof76) to 50.33 kD (BnaDof33). The GRAVY (Grand Average of Hydropathy) values of the BnaDofs reflected the hydrophilic nature of these proteins.

We also identified 62 *B. oleracea* genes as members of the Dof TF family (Table S2). It is worth to note that in addition to the 117 *BnaDof* genes, one other gene (*BnaA07g02590D*) showed the presence of the Dof domain along with a Syntaxin domain. Further investigation revealed that this gene was homologous to *Bra002057*, which is reported as a *B. rapa* Dof gene family member (Ma et al. 2015). However, this gene is orthologous to Arabidopsis *SYNTAXIN OF PLANTS 21* (*SYP21, At5g16830*). Thus, we decided to exclude *BnaA07g02590D* and *Bra002057* from our analysis as these genes can be classified as members of the SNARE (SNAP Receptor) protein family (Lipka et al. 2007). Additionally, while performing the phylogenetic analysis, these genes failed to produce bootstrap replications.

### Phylogenetic relationships of the *Dof* gene family in *B. napus*

We explored the phylogenetic relationships between the *Dof* families in *B. napus* and Arabidopsis by first performing alignment of the 117 *BnaDofs* and 36 Arabidopsis *Dof*s using CLUSTALW (Larkin et al. 2007). Then the multiple sequence alignment was used to construct the unrooted phylogenetic tree in MEGA7.0 (Kumar et al. 2008) using the neighbour-joining method with 1000 bootstrap values (Figure 2). We then classified *BnaDofs* into four major groups: A, B, C and D and the following nine subgroups: A, B1, B2, C1, C2.1, C2.2, C3, D1, D2, based on the phylogenetic tree. The major group was C, with 35% of *BnaDofs* followed by group D with 28% BnaDofs. Among the subgroups, D2 was the largest, comprising of ~18% *BnaDofs*. Least number of BnaDofs belonged to subgroup C1.

**Figure 2.**
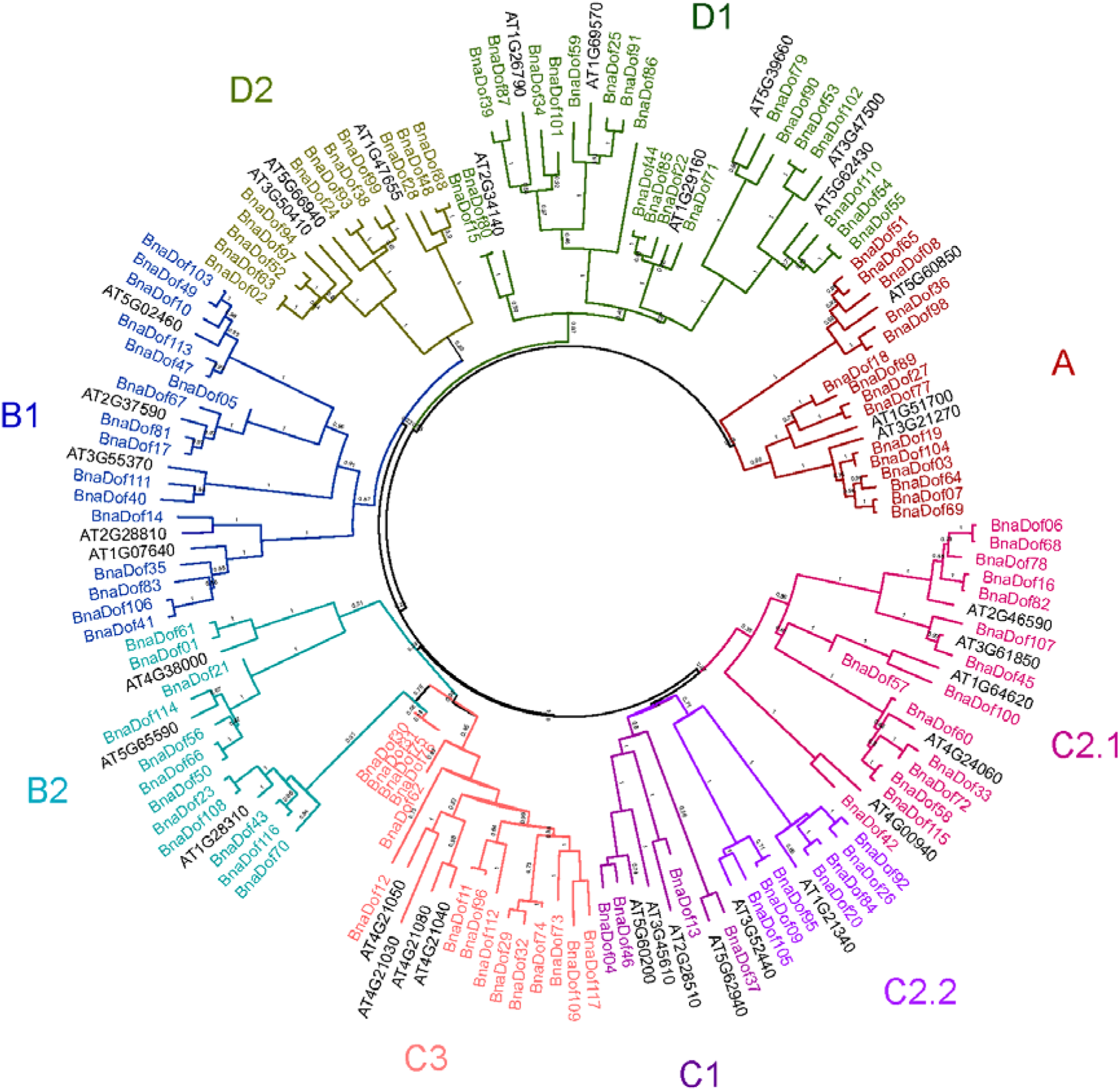
Phylogenetic tree of *B. napus* and Arabidopsis Dof proteins. The full-length amino acid sequences were aligned using CLUSTALW, and the phylogenetic tree using the Neighbor-Joining method was constructed using MEGA7. The percentage of replicate trees in which the associated taxa clustered together in the bootstrap test (1000 replicates) are shown next to the branches. The evolutionary distances were computed using the Poisson correction method. The analysis involved 153 amino acid sequences.

We further explored the phylogenetic relationship between the Dof gene family in *B. napus, B. oleracea, B. rapa* and Arabidopsis. As mentioned earlier, there are 76 (75 included in our analysis) *B. rapa Dofs* and 36 Arabidopsis *Dofs* and we identified 117 and 62 *Dof* genes in *B. napus* and *B. oleracea*, respectively (Ma et al. 2015; Yanagisawa 2002). So, a total of 290 Dof protein sequences were utilised to construct the phylogenetic tree. Based on the resulting phylogenetic tree, the *Dof* gene family can be classified into four major groups and nine subgroups as described earlier (Figure S1). So, a total of 290 Dof protein sequences were utilised to construct the phylogenetic tree. Based on the resulting phylogenetic tree, the *Dof* gene family can be classified into four major groups and nine subgroups, as described earlier (Figure S1). The tree illustrates the expansion and divergence of the *Dof* gene family from Arabidopsis to *B. napus*. Few homologues from the two diploid progenitors were lost in *B. napus* during hybridisation, and few underwent further duplications after hybridisation. However, the majority of *BnaDofs* were consistently inherited from their progenitors (Table S3). Based on the known chromosomal locations we confirmed a total 43 and 33 gene pairs which maintained their relative positions between the *B. rapa’* genome and A_n_ sub-genome in *B. napus* and *B. oleracea’* genome and C_n_ sub-genome in *B. napus*.

### Gene structure and conserved motifs of BnaDofs

To gain further understanding into the structural diversity of *BnaDofs*, we studied exon-intron organisation and identified the presence of conserved protein motifs (Figure 3). The analysis revealed 61 *BnaDofs* with no introns, whereas 46, 8 and 2 *BnaDofs* had one, two and three introns, respectively. Majority of the genes within a given subgroup showed similar exon-intron organisation. For example, all the C1 subgroup *BnaDofs* had one intron, and most members of subgroup D2, A, B2, C3 and C2.2 had no introns. Similarly, *BnaDofs* belonging to subgroup B1 had at least one intron. Most diverse gene structure organisation was observed in the members of subgroup D2, ranging from zero to three introns.

**Figure 3.**
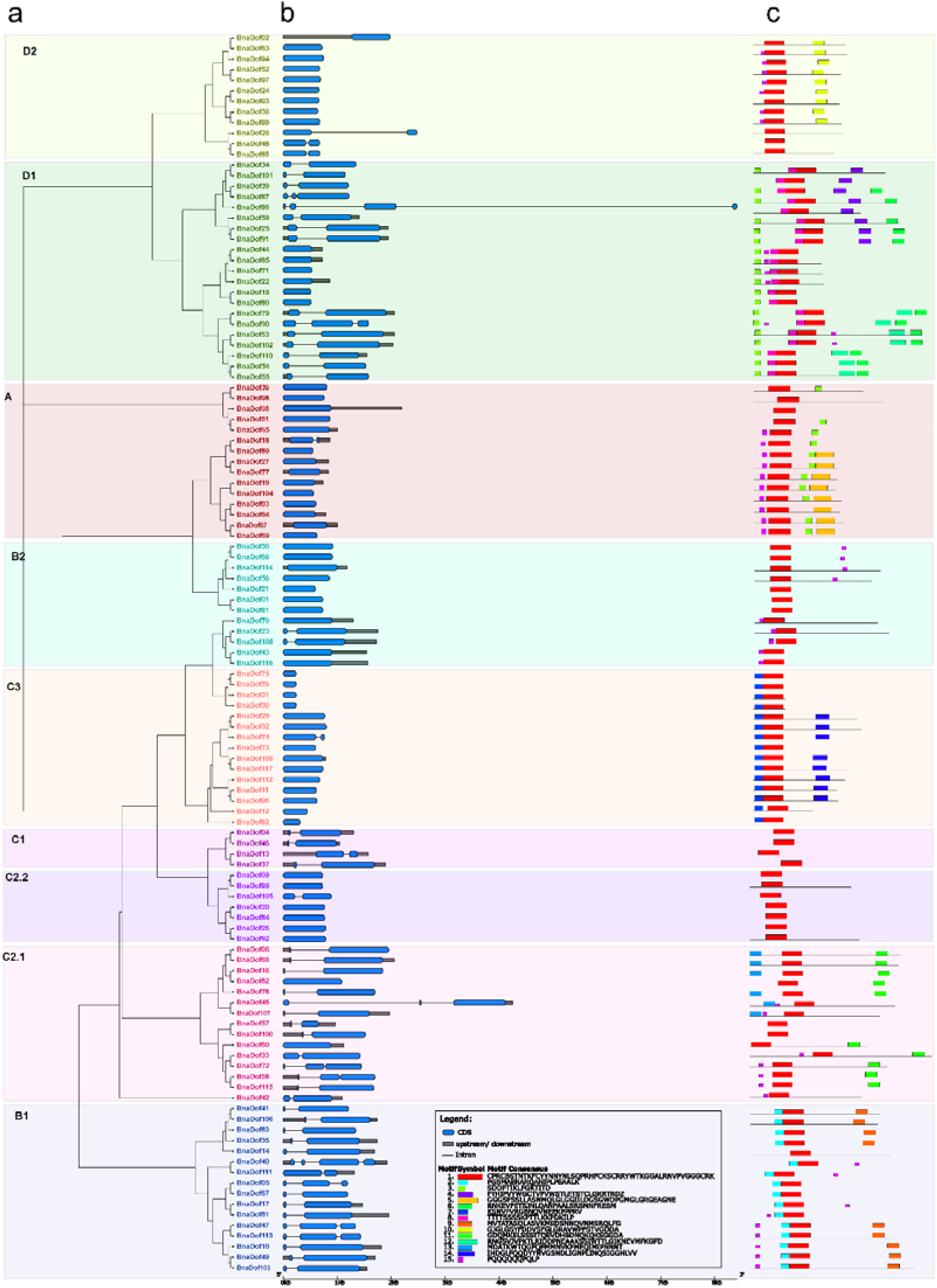
Phylogenetic relationships, gene structure, and conserved protein motifs in the members of *Dof* gene family in *B. napus* (a) Phylogenetic relationships of BnaDofs. The phylogenetic tree was constructed with MEGA 7.0 using the neighbor-joining (NJ) method with 1000 bootstrap replicates and Poisson correction method. The nine Dof groups are displayed in different text colour and enclosed in the respective colour boxes; **(b) Gene structure of *BnaDof* genes**. Grey boxes indicate untranslated 5’- and 3’-regions; blue boxes indicate exons; black lines indicate introns. Scale bar represents gene length; **(c) Distribution of conserved motifs in BnaDof proteins.** The sequence of each motif (1-15) displayed in different coloured boxes is provided in the legend.

Motif analysis performed using MEME showed the presence of a highly conserved motif-Motif 1, which represents the Dof-type domain, across all the 117 BnaDofs’ amino acid sequences (Bailey et al. 2015). The conserved nature of motif distribution within the BnaDofs’ protein subgroup also highlighted their phylogenetic relationships. It is worth mentioning that conserved motif distribution also existed among members of different clades within a subgroup. For example, BnaDof28, BnaDof48 and BnaDof88 which belong to one clade in subgroup D2 show the presence of only one out of three conserved motifs that are found in the other D2 subgroup members. Thus, the results suggest that gene and protein structural divergence across subgroups probably governs the functional diversity in the Dof subgroups.

### Orthologous gene clustering of Dof gene family in B. napus, B. oleracea, B. rapa and Arabidopsis

To understand the evolutionary relationship of Dof gene family among the four important members of the Brassicaceae family - *B. napus, B. oleracea, B. rapa* and Arabidopsis --, we carried out an orthology analysis in OrthoVenn2 web platform (Xu et al. 2019). The identified orthologous clusters in the four species are illustrated in Figure 4. 135 Dof proteins from all four species were clustered in 29 orthologous groups. We also identified 25 *Brassica* specific clusters with 88 Dof proteins from the three *Brassica* species. One Arabidopsis Dof protein (AT3G45610), 17 BnaDofs and 6 *B. rapa* Dofs did not cluster in any orthologous group and were identified as singletons. No singletons were identified in *B. oleracea*. Further, there were 7 clusters (17 Dofs) between *B. rapa* and *B. napus*, 7 clusters (12 Dofs) between *B. oleracea* and *B. napus* and 2 clusters between *B. rapa* and *B. oleracea*. The absence of *B. napus’* gene in *B. rapa - B. oleracea* specific clusters suggest that genes belonging to these clusters might have been lost during the hybridisation event. A detailed list of orthologous gene clusters and singletons is provided in Table S4a-b.

**Figure 4.**
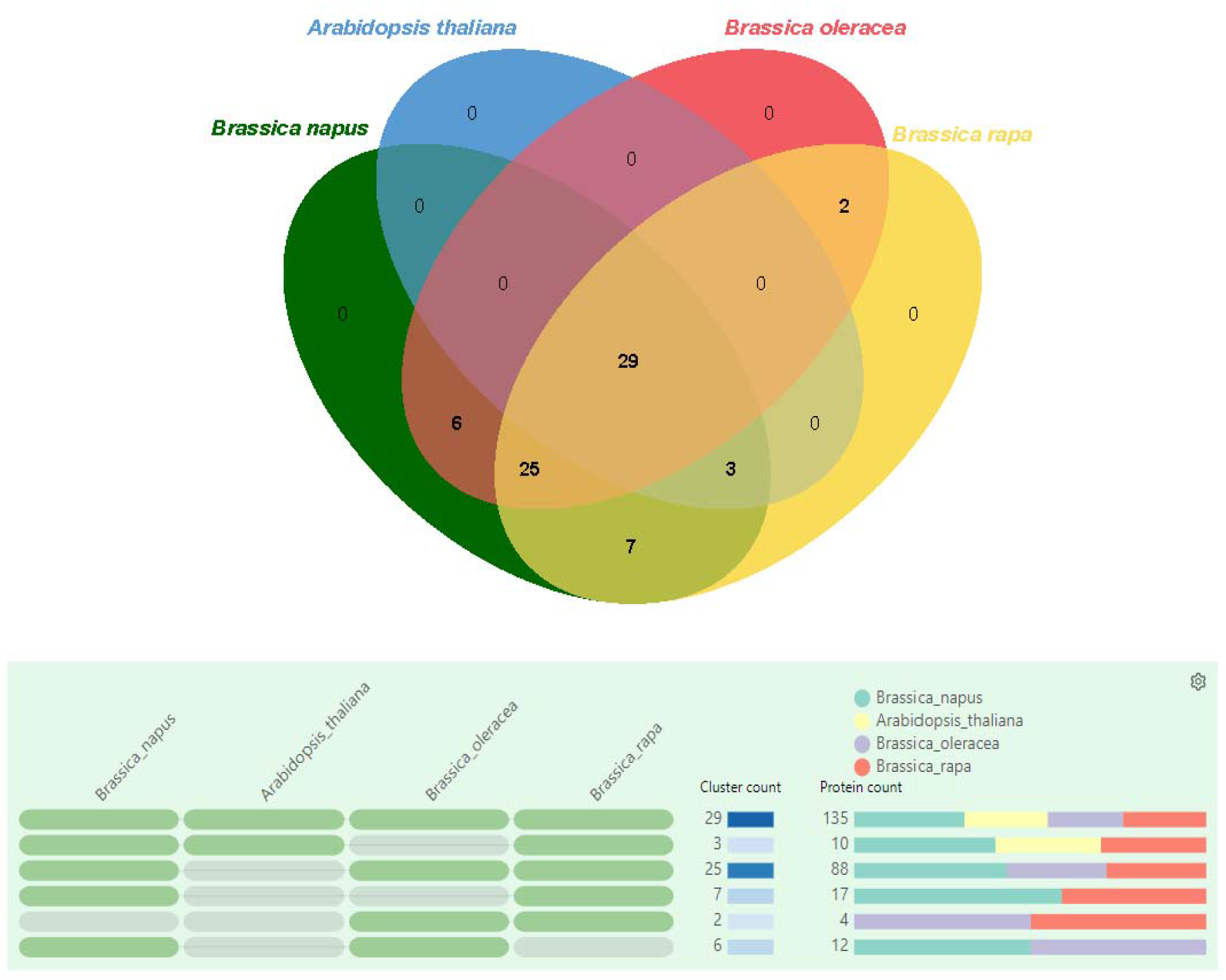
Orthologous gene clustering analysis. The orthologous gene clusters between the Dof gene family in *B. napus, B. oleracea, B. rapa* and Arabidopsis, were identified and visualised using the OrthoVenn2 web platform. The e-value cut off 1e-10 was used for the analysis.

### Evolution and Divergence of BnaDofs

The expansion of a gene family occurs because of duplication events arising at a whole-genome level or a small scale. We first identified duplicated gene pairs among *BnaDofs* based on the sequence similarity (Table S5a). We found 128 gene pairs with >80% sequence similarity. Gene pairs are identified as tandemly duplicated if the new gene/sequence is found adjacent (within 100kb window) to the duplicated genomic region. Based on these criteria, three *BnaDof* gene pairs were identified as tandemly duplicated (*BnaDof30-31, BnaDof 75-76, BnaDof 86-87*). Rest of the *BnaDofs* gene pairs underwent interspersed duplications. Furthermore, BLASTP and MCScanX based methods identified 85 segmental, 26 dispersed, 4 tandem and 1 proximal duplication event (Table S5b) (Korf et al. 2003; Wang et al. 2012). These results highlight that segmental duplication events played a critical role in shaping the *Dof* gene family in *B. napus*.

Nucleotide substitutions that producing an amino acid change are termed non-synonymous, and those that do not are termed synonymous. The ratio of non-synonymous to synonymous substitutions (Ka/Ks) in a protein-coding gene reflects the magnitude and direction of selection pressure acting on a protein sequence (Yang and Bielawski 2000). Ka/Ks value <1 indicates that a gene pair has experienced negative or purifying selection (acting against change), whereas Ka/Ks >1 indicates positive or adaptive selection (driving change) and Ka/Ks = indicates neutral selection (Li et al. 2009). Thus, we calculated the Ka/Ks ratio among the duplicated gene pairs (Table S5a). Among the 128 identified duplicated *BnaDof* gene pairs, six gene pairs underwent neutral selection, and the remaining underwent negative or purifying selection (Ka/Ks<1).

The syntenic relationships between chromosome segments of different species can provide valuable insights into the origin of the gene family members. For the synteny analysis, only the genes with known chromosomal locations were considered. We performed a syntenic analysis between Dof genes from *B. napus* and Arabidopsis (Figure 5a). In accordance with our orthology analysis, we identified orthologs for 35 Arabidopsis genes in *B. napus*. In addition, we also constructed a synteny map between Dof genes from *B. napus, B. oleracea* and *B. rapa*. 58 out of 117 genes in *B. napus*, and 2 out of 62 were discarded due to uncertain chromosomal locations (Figure 5b). Out of the remaining 59 *Bna*Dofs, 98.3% of genes were placed in collinear blocks. In *B. rapa*, 56.6% (43/75 Dof genes) and in *B. oleracea*, 78.3% (47/60 Dof genes) were placed in collinear blocks. Since the same gene in *B. napus* could be in a colinear block relative to *B. rapa* but not relative to *B. oleracea*, the numbers reported for *B. rapa* and *B. oleracea* indicates how many genes are collinear with the corresponding orthologs in *B. napus*. For *B. napus*, instead, this number report the number of genes that are collinear in at least one of the other species.

**Figure 5.**
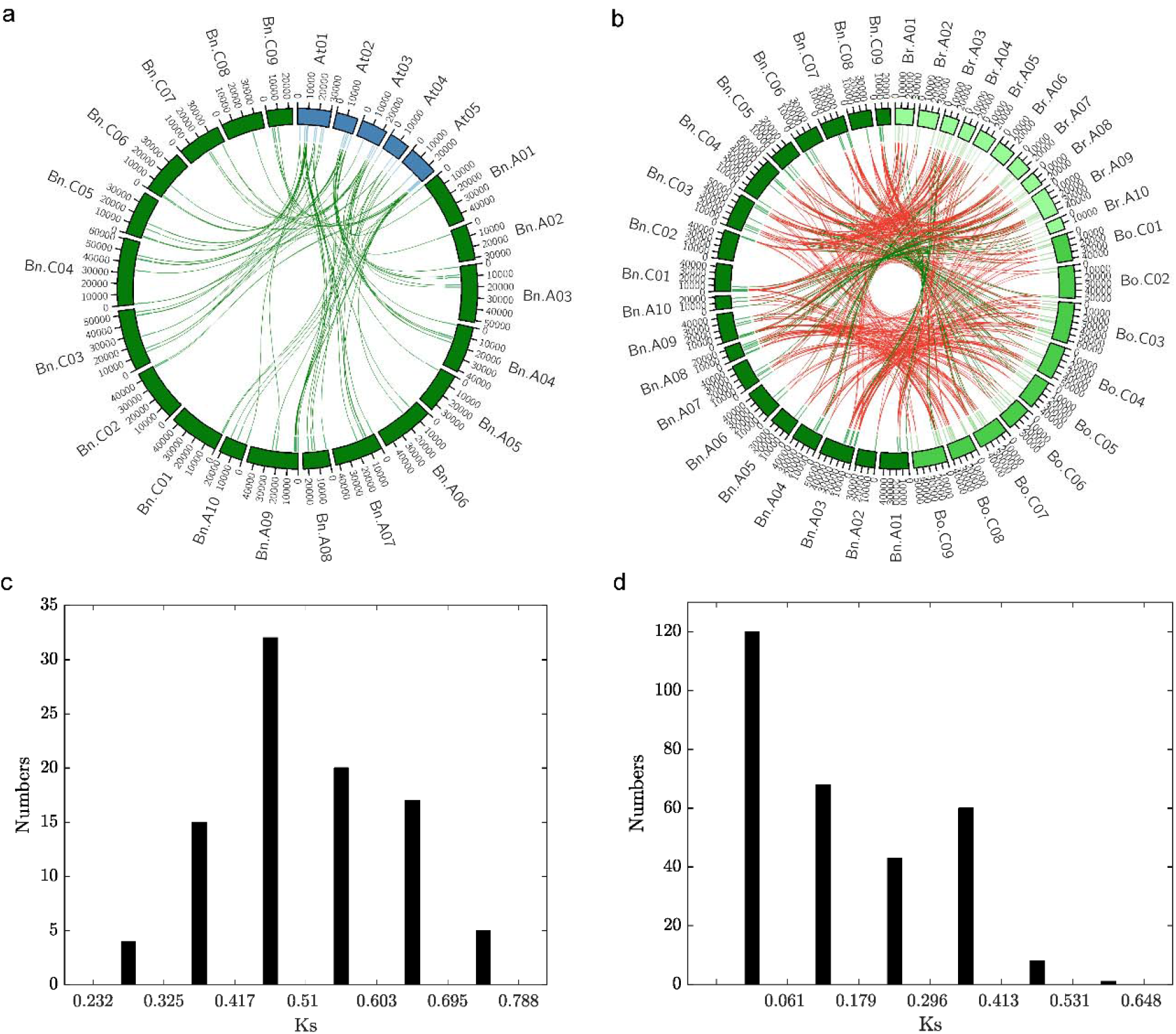
Synteny analysis (a) Synteny of Dof gene family between A. thaliana and B. napus. Ideograms of chromosomes of *A. thaliana* (blue) and *B. napus* (dark green) are displayed in the outer circle, while in the inner circle indicates the positions of the genes in the corresponding chromosomes. Green links connect orthologous genes between the two species. The chromosome scale is in Kb. At: *Arabidopsis thaliana;* Bn: *Brassica napus;* **(b) Synteny of Dof transcription factors within the *Brassicaceae* genus.** Ideograms of chromosomes of *B. rapa* (pale green), *B. oleracea* (lime green) and *B. napus* (dark green) are displayed in the outer circle, while in the inner circle indicates the positions of the genes in the corresponding chromosomes. Green links connect collinear orthologous genes, while red links connect orthologues genes that underwent chromosomal translocation events. The chromosome scale is in Kb. Bn: *Brassica napus;* Bo: *Brassica oleracea;* Br: *Brassica rapa;* **(c) Density of Ks values Dof orthologous gene pairs between *B. napus* and Arabidopsis**. Analyses were conducted using the Nei-Gojobori model in MEGA7.0. All ambiguous positions were removed for each sequence pair; **(d) Density of Ks values of Dof orthologous gene pairs between *B. napus, B. oleracea* and *B. rapa***. Analyses were conducted using the Nei-Gojobori model in MEGA7.0. All ambiguous positions were removed for each sequence pair.

Furthermore, we estimated the divergence time of the Dof gene family between Arabidopsis and *B. napus* by calculating the Ks values of the identified orthologous gene pairs (Table S6a). The Ks value for all the orthologous pairs ranged from 0.28 to 0.74, with an average of 0.51 (Figure 5c). Using the estimate of mutational rate, R = 1.5 × 10^-8^ synonymous substitutions per site per year (Koch et al. 1999; Koch et al. 2001), the average estimated divergence time of the Arabidopsis and *B. napus Dof* gene family was ~17Mya. Our results agree with the reported estimated divergence time (14-24 MYA) of the Arabidopsis and *B. napus* lineage (Cheung et al. 2009; Koch et al. 2000). We also calculated the Ks values of orthologous gene pairs between the three Brassica species (Table S6b). The Ks value ranged from 0.0024 to 0.5896. with an average of 0.15 (Figure 5d). The average divergence time was ~5Mya (80,000 years - 19.5Mya). The hybridisation event between *B. rapa* and *B. oleracea* took place around 7500-12,500 years ago, and the Brassica whole-genome triplication event is estimated to have taken place around 9-15Mya (Chalhoub et al. 2014; Cheng et al. 2014). Overall, these results indicate the divergence of Brassica genus from *Arabidopsis*, followed by whole genome triplication, hybridisation of *B. rapa* and *B. oleracea* to form *B. napus*, as well as gene loss and rearrangements shaped the *Dof* gene family in *B. napus*.

### Functional annotation of BnaDofs and Promoter analysis

Functional annotation allows detailed evaluation of proteins with unidentified molecular function, biological process, or cellular component. In the cellular component gene ontology category, all the BnaDofs were associated with “nucleus”, and the majority of them were associated with “integral component of membrane” terms. A Dof gene family is a transcription factor family so, it was expected that in the molecular process category the BnaDofs would be associated with terms such as “DNA binding” and “DNA-binding transcriptional factor activity”. In the biological process category, BnaDofs was not only associated with “regulation of transcription” but also with a number of other terms related to organ development, vegetative to reproductive transition, light signalling, response to different hormones, cell differentiation, oxidation-reduction among others. A detailed summary of functional annotation results along with descriptors is provided in Table S7.

To gain further understanding of the functional roles of *BnaDofs*, we used the PlantCARE database to identify potential cis-regulatory elements present upstream of the coding regions (1.5kb upstream) (Lescot et al. 2002). Several cis-acting regulatory elements were found in the promoter region of *BnaDofs*, and we classified them into three categories: developmental, stress-responsive, and hormone-responsive (Figure 6). Among the development related *cis*-elements, we identified elements regulating light responsiveness (G-box, Box-4, GT1-motif, 3-AF1, AAAC-motif, Sp1, MRE), circadian rhythm (circadian), meristem expression (CAT-box) and differentiation of palisade mesophyll cells (HD-Zip-1). *cis*-acting regulatory elements related to light responsiveness, especially the G-box, Box-4, and GT1-motif elements were present in ~66%, ~77%, and ~50% *BnaDofs*. CCGTCC-box, which is a development-related cis-element, was also found in the promoters of 11 *BnaDofs*, out of which 7 *BnaDofs* belonged to the D major group.

**Figure 6.**
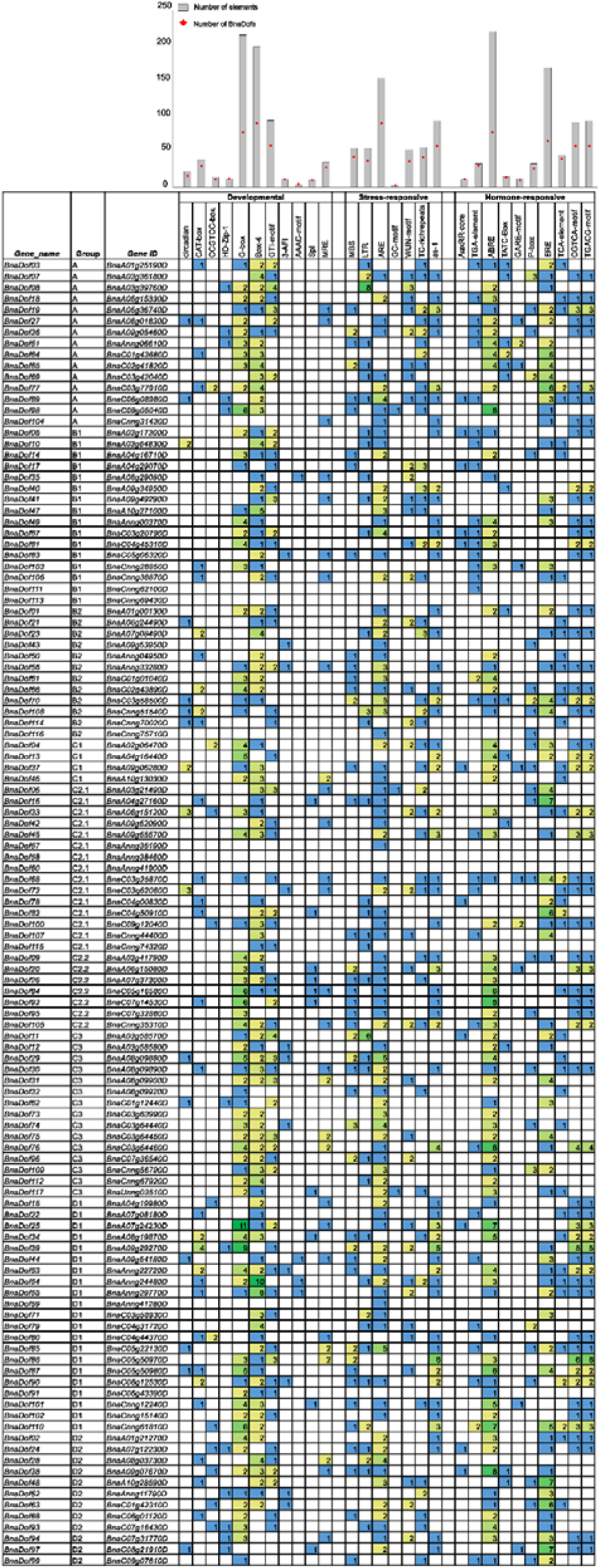
*cis*-acting regulatory elements in the promoter region of *BnaDof* genes. The cis-acting regulatory element analysis in the promoter region (1.5 kb upstream of translation initiation site) of *BnaDof* genes was performed using the PlantCARE database. The number of each cis-acting element in the promoter regions of *BnaDof* genes are represented in three major categories: developmental, stress-responsive, and hormone-responsive. On top, the bar graph represents the total number of each cis-acting element present in *BnaDofs* (Grey box) and the corresponding number of *BnaDofs* promoters carrying a particular cis-element (Red diamond). The details of the ciselements are provided in Table S10.

Further, we detected the presence of stress-responsive cis-elements MBS (involved in drought inducibility), LTR (low-temperature responsive), WUN-motif (wound responsive), TC-rich repeats (defence and stress-related), ARE (anaerobic induction), and GC-motif (anoxic specific inducibility) in 43, 37, 36, 42, 90 and 3 *BnaDof* promoters, respectively. As-1 cis-element reported to be present in pathogenesis-related genes in plants was also detected in the promoters of 58 *BnaDofs*. Stress signalling and hormone signalling operate at an intertwined level in regulation of plant stress-responsive gene expression. Therefore, we identified hormone-responsive cis-elements in the *BnaDof* promoters. Among all the hormone-responsive elements, 219 ABRE elements were present in 77 (68%) *BnaDofs*. Cis-elements responsive to auxin (AuxRR-core, TGA-element), gibberellin (TATC-box, GARE-motif, P-box), salicylic acid (TCA-element), ethylene (ERE) and methyl jasmonate (CGTCA and TGACG) were also present in the *BnaDof* promoters. In addition to the above-mentioned cis elements, we encountered several other cis-acting elements such as the AE-box (part of a module for light response), MYB, Myb-binding site, MYC, TATA-box and CAAT box which was present in the *BnaDof* promoters.

### Tissue-specific and abiotic stress responsive expression profiling of *BnaDofs*

To analyse the tissue-specific expression patters of *BnaDofs*, four tissues were compared: young root, stem, leaf, and flower buds (Table S8a). The distinctive tissue-specific expression of *BnaDofs* can be grouped into eight (T.I - T.VIII) clusters, as illustrated in Figure 7a. Majority of the *BnaDofs* belonging to cluster T.II show higher expression in young roots. Similarly, T.III and T.VIII show higher expression in flower buds, and stem, respectively. Additionally, *BnaDofs* belonging to cluster T.I, T.IV, T.V and T.VII show higher expression in at least two tissues. Interestingly, in cluster T.VI, *BnaDof01, BnaDof34, BnaDof65* and *BnaDof 101* have higher expression in leaves and flower buds, *BnaDof61, BnaDof62, BnaDof66, BnaDof101* and *BnaDof116* show higher expression in leaves whereas the expression of *BnaDof11, BnaDof12, BnaDof29, BnaDof32, BnaDof73* and *BnaDof96* was undetectable in any of the four tissues. Preferential expression within the tissue-specific cohort of *Dof* family group was also noticeable. For instance, *BnaDof* members belonging to group C2.1 showed higher expression in young roots and stem, more than 50% of the members of A and B1 *Dof* group showed preferential expression in stem and young roots, respectively.

Brassica Expression DataBase (BrassicaEDB)-A gene expression database for Brassica crops (version 1.0) was released recently (Chao et al. 2020). To get a comprehensive understanding of developmental expression patterns of BnaDofs, we compiled the expression maps of BnaDofs, across different tissues and developmental stages as available on Brassica EDB (Supplementary Folder). The expression levels of *BnaDof11, BnaDof12, BnaDof29, BnaDof32, BnaDof73* and *BnaDof96* were undetectable in young roots, stem, leaves, and flower buds based on our RNA-Seq analysis (Figure 7a). Thus, we observed the expression maps of these genes on BrassicaEDB. *BnaDof73* showed limited expression in cotyledons (48 hours after germination), seed-13 days after fertilisation and in the seed coat (inner integument). Similarly, *BnaDof96* and *BnaDof11* were only expressed in the seed coat (inner integument), *BnaDof32* and *BnaDof29* in seeds-13 days after fertilisation, and *BnaDof12* in seeds-10 days after fertilisation.

**Figure 7.**
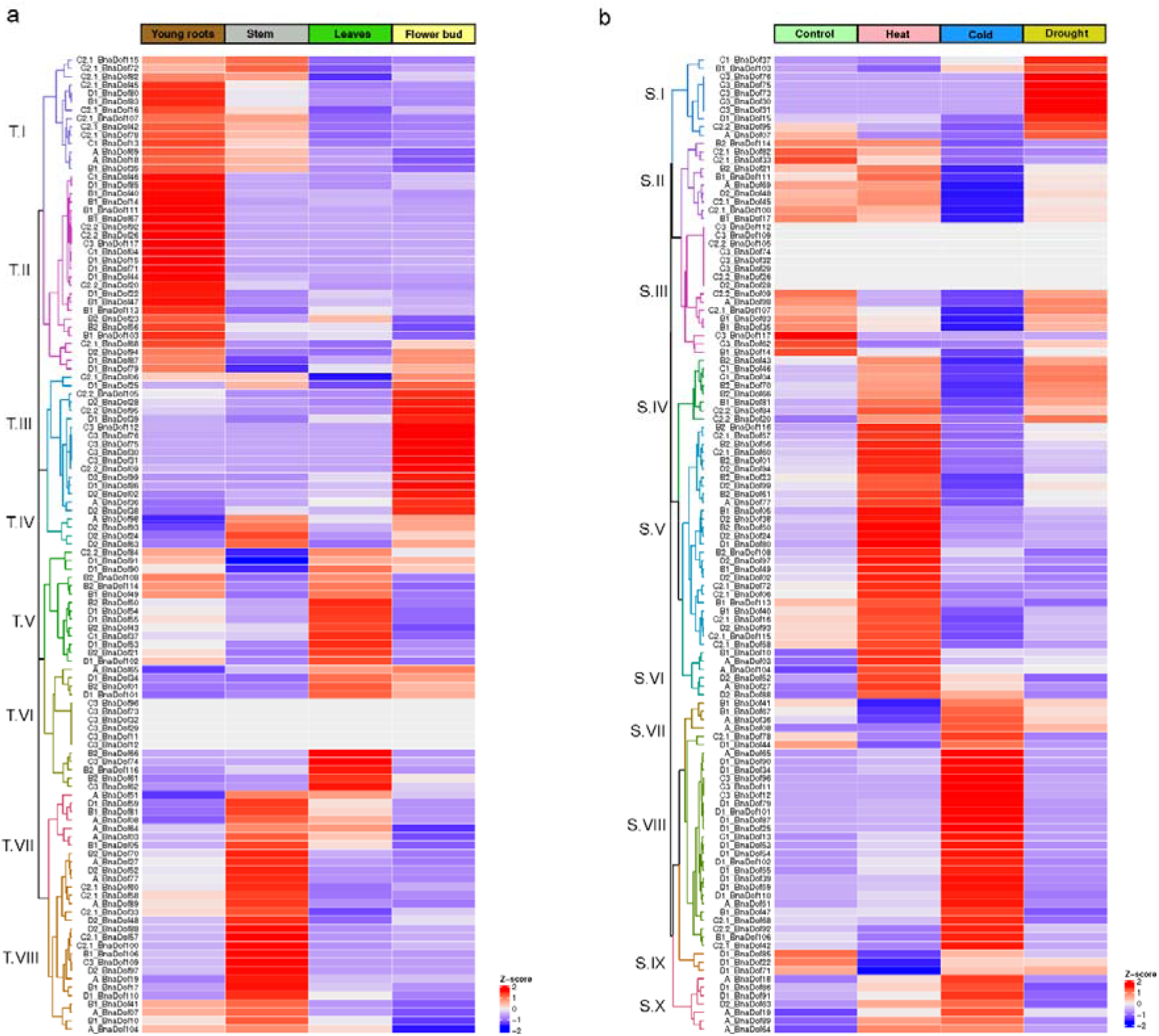
(a) Heat map representation of the tissue-specific expression of BnaDofs. RNA-seq data were obtained from root, stem, leaf, and flower of *B. napus;* (b) **Heat map representation of the stress-responsive expression of BnaDofs**. RNA-seq data were obtained from three weeks old B. napus seedlings grown at control conditions (16/8 h photoperiods at 22 °C, RH 50%, and light intensity of 230-240 μmol m^-2^ s^-1^) and exposed to Heat: 35°C for 24h, Cold: 4°C for 24h, and Drought: 22°C, 25% PEG□6000 for 24h. Hierarchical clustering of BnaDof expression profiles was performed using the Euclidean distance method and complete clustering method. The scale bar represents the Z-score (scaled TPM values).

We further investigated the changes in expression patterns of *BnaDofs* in three-week-old *B. napus* seedlings exposed to various abiotic stresses. Figure 7b illustrates the expression patterns of *BnaDofs* in control conditions and upon exposure to heat stress, cold stress, and drought conditions (Table S8b). The expression profiles of *BnaDofs* were grouped into ten clusters (S.1 - S.X). Numerous *BnaDof* genes showed changes in expression profiles upon exposure to extreme temperature. *BnaDofs* grouped in clusters S.VII, S.VIII, and S.X were upregulated in response to low-temperature stress. Members of the *BnaDof* gene family showed, especially upon exposure to cold stress. Cluster S.IV, S.V, and S.VI genes were up-regulated upon exposure to heat stress. Majority of the heat-responsive *BnaDofs* were upregulated. Interestingly a fraction of BnaDofs which were upregulated in response to heat stress showed downregulation in response to cold stress and vice versa. For example, BnaDofs clustered in S.IV were upregulated in response to heat and down-regulated in response to cold, and an opposite trend was seen in genes clustering in S.VII. Genes belonging to cluster S.I, and S.IV showed significant upregulation under drought conditions and S.II cluster *BnaDofs* were downregulated. However, the majority of *BnaDofs* were not drought-responsive. The expression of some *BnaDofs* belonging to cluster S.III *(BnaDof112, BnaDof109, BnaDof105, BnaDof74, BnaDof32, BnaDof29, BnaDof26* and *BnaDof28*) was undetectable or did not show any changes in expression upon abiotic stress exposure. This might be due to no detectable expression of these genes in three-week-old seedlings and probably indicate these genes might not be involved in heat, cold and drought stress response.

## Discussion

*Brassica napus*, the second largest economically important oilseed crop is an allopolyploid formed as a result of spontaneous pairwise hybridisation of *B. rapa* and *B. oleracea*. The availability of the *B. napus* genome provides opportunities for identification of important gene families. One of the important plant-specific transcription factor family is Dof zinc finger gene family. This gene family has been described in several plant species including Arabidopsis, rice, wheat, tomato, pepper, Chinese cabbage, and cucumber, among others but not in *B. napus* (Lijavetzky et al. 2003; Liu et al. 2020; Ma et al. 2015; Wen et al. 2016; Wu et al. 2016; Yanagisawa 2002). In this study, we performed *in-silico* genome-wide identification, comparative, and evolutionary analysis of the *Dof* gene family in *B. napus*. Here, we report 117 genes as putative members of the *Dof* gene family in *B. napus* which is the largest number of *Dof* genes ever reported in eudicots.

### Systematic analysis of *BnaDofs*

According to the phylogenetic analysis of *Dof* genes in Arabidopsis and rice reported by Lijavetzky et al. (2003), members of *Dof* gene family are classified into nine groups. The 117 *BnaDofs* identified in our study were also classified into nine groups based on the phylogenetic analysis between the identified *BnaDofs* and Arabidopsis *Dof* genes. We further performed a phylogenetic analysis between Arabidopsis, *B. napus, B. rapa* and *B. oleracea*. Our present study also identified 62 *Dof* genes in *B. oleracea*. In *B. rapa*, 76 *Dof* genes are reported (Ma et al. 2015; Yanagisawa 2002). However, we included only 75 *B. rapa* Dof in our analysis since one gene contained a Syntaxin domain along with the Dof domain and was orthologous to a *SYNTAXIN OF PLANT 21* gene in Arabidopsis. The *Dof* gene family across the *Brassica* species also clustered phylogenetically in nine groups. Further, comparative analysis of the gene structure and conserved protein domains highlighted conserved exon-intron organisations and distribution of protein motifs followed by the members of a group. The Dof domain (motif 1, Figure 3c) was conserved across all BnaDofs, and similar motif distribution pattern can be seen across the members belonging to the same group. Furthermore, the structural conservation of *BnaDofs* was in accordance with Dof genes reported in other plants such as Arabidopsis, rice, and *B. rapa* (Lijavetzky et al. 2003; Ma et al. 2015; Noguero et al. 2013).

### Expansion and divergence of DOF gene family in *B. napus*

Genome wide studies identifying gene families in *B. napus* have reported frequent expansion of gene families such as HSF, GST, CRF, TLP, bHLH to name a few (Lohani et al. 2019; Miao et al. 2020; Wang et al. 2020a; Wang et al. 2020b; Wei et al. 2019). The difference in the size of *Dof* gene family from Arabidopsis to *B. napus* indicates the expansion of the *B. napus Dof* gene family. We performed orthologous gene clustering, synteny analysis and molecular evolutionary analysis to understand the expansion of the BnaDofs. The average synonymous base substitution rate between *B. napus* Dof genes and their Arabidopsis orthologs was calculated as 0.51 (0.28 to 0.74). We further estimated the divergence time of ~17Mya, and it was constant with the time Arabidopsis and *Brassica* lineages diverged i.e., 14-24 Mya (Koch et al. 2000). Similarly, the calculated maxima, minima, and average Ks values of the orthologous gene pairs between the *Brassica* species highlights that the whole genome triplication events (9-15 Mya) and the hybridisation event (7500 years ago) and led to the expansion of *Dof* gene family in *B. napus* (Chalhoub et al. 2014; Cheung et al. 2009). Expansion of gene families due to whole genome and local gene duplication events might be an effective strategy in plants for adapting to the ever-changing environmental conditions.

### Distinct expression patterns of *BnaDofs* during development

Tempo-spatial expression profiles of *BnaDofs* in association with functional annotations and cis-element analysis of BnaDof promoters indicate their preferential expression in different tissues suggesting diversification of function during organ development. The functional characterisation of Dof proteins in different plants has indicated their association with light-responsiveness, phytochrome signalling, seed germination, tissue-specific expression in endosperms, vascular tissue development, leaves or guard cells (Le Hir and Bellini 2013; Noguero et al. 2013; Ruta et al. 2020). Here in this section, to provide better clarity, we will be discussing the *BnaDof* genes in terms of orthologous relationships with Arabidopsis *Dof* genes rather than in terms of classified groups as different schemes for *Dof* gene family classification are available in the literature (Le Hir and Bellini 2013; Lijavetzky et al. 2003; Yanagisawa 2002).

Light is an indispensable environmental cue which regulates developmental processes such as photomorphogenesis, seed germination, flowering, and several other metabolic and cellular processes (Legris et al. 2017). In plants, the first *Dof* TF reported in maize was shown to be involved in light signalling (Yanagisawa 1995; Yanagisawa 2000; Yanagisawa and Sheen 1998). The *BnaDof* promoters are enriched for the presence of light-responsive cis-acting elements. Several *Dof* genes in plants are associated with the regulation of photomorphogenesis, seed germination and development (Noguero et al. 2013; Ruta et al. 2020).

Circadian rhythms occur in plants to respond to daily and seasonal changes and synchronise their developmental programme based on the day length (Inoue et al. 2018). In addition to light-responsive cis-elements, few BnaDof promoters also showed the presence of circadian clock associated cis elements (Figure 6). In Arabidopsis, Cycling DOF factors (*CDF1, CDF2, CDF3* and *CDF5*) are reported to regulate photoperiodic flowering response (Fornara et al. 2009; Song et al. 2013). In Arabidopsis, *CDF1* represses the transcription of a core circadian clock signalling gene *CONSTANS (CO)* (Fornara et al. 2009; Imaizumi et al. 2005). *CO* gene is involved in regulation of flowering under long days and its repression by *CDF1* represses flowering in Arabidopsis (Fornara et al. 2009; Shim et al. 2017). Its ortholog in *B. napus, BnCDF1* (*BnaDof54* in our study) has also been reported to play role in flowering (Xu and Dai 2016). *BnaDof54*, functionally annotated to be involved in flower development shows very high expression in flower. Similarly, *BnaDof55*, which is also orthologous to *CDF1*, was functionally annotated to be related to flower development and showed higher expression in flower indicating a similar functional role.

Light is essential for the conversion of the inactive Pr form of phytochrome into the active Pfr, which then activates the process of seed germination (Shinomura et al. 1994). In Arabidopsis, *DAG1, DAG2, COG1* and *OBP3* are reported to regulate in seed germination and hypocotyl elongation (Gualberti et al. 2002; Papi et al. 2000; Park et al. 2003; Ward et al. 2005). In response to light, *DAG1* represses seed germination and *DAG2* activates seed germination, thus acting antagonistically (Gualberti et al. 2002). In Arabidopsis, the expression of *DAG1* and *DAG2* is detected in vascular tissues but not in seed suggesting regulation of long-distance light related signalling pathways. *B. napus* genes orthologous to *DAG1* is *BnaDof45* and *DAG2* are *BnaDof06* and *BnaDof68*. These genes also showed comparatively higher expression in tissues other than seed or embryo.

The first reported Dof protein to regulate phytochrome mediated signalling involved in seedling development was *COG1* (Park et al. 2003).]. *COG1* interacts with *Phytochrome Interacting Factors (PIF4* and *PIF5*), activates Brassinosteroids biosynthesis and promotes hypocotyl elongation (Wei et al. 2017). It is also reported that *COG1* controls the expression of *PRX2* and *PRX25*, which are associated with seed longevity and thereby regulates seed tolerance (Renard et al. 2020). Four *BnaDof22, BnaDof44, BnaDof71* and *BnaDof85* were identified as orthologous to *COG1(At1g29160)* and were functionally annotated to be associated with seed coat development and showed very high to moderate expression in seed coat. Arabidopsis’ *OBP3* gene has been shown to be involved in the repression of hypocotyl elongation in a light-dependent manner (Ward et al. 2005). Among the *BnaDofs* one of the orthologs of *OBP3, BnaDof40* showed very high expression in hypocotyl, suggesting similar function.

Arabidopsis *Dof* gene, *At3g45610* also known as *Dof6* or *Dof3.2* to negatively regulates seed germination (Rueda-Romero et al. 2012). Orthology analysis revealed the absence of *Dof6* orthologous gene across *B. napus Dof* gene family. Gene loss due to evolution and divergence of *BnaDof* genes might have either resulted in a loss of function or neofunctionalization. It is worth mentioning that the cis-element analysis of the *BnaDof* promoters also revealed the presence of cis-elements which are related to or act as binding sites for other transcription factors (MYB, MYC, ARF, MADS-boxes). Similar diversification of binding sites is reported for Arabidopsis *Dof* promoters, indicating potential relationships between *Dof* TFs and other TFs in regulating diverse plant developmental processes (Yilmaz et al. 2010).

### Potential role of *BnaDofs* in abiotic stress response

*Dof* TFs have also been reported to participate in response to abiotic stress response (Waqas et al. 2020). We investigated the expression profiles of *BnaDofs* in response to heat, cold and drought stress. Our analysis highlights the temperature responsiveness of *BnaDofs* with a majority of *BnaDofs* showing differential regulation upon exposure to low temperatures. Under the high-temperature majority of the differentially regulated *BnaDofs* were upregulated. A recent study in *B. napus* exploring the cold-responsive TFs reported the changes in expression of *Dof* genes in response to cold stress response and suggested their possible role in imparting cold tolerance (Ke et al. 2020). It is important to note that depending upon the variety and even plant species, the expression profiles of *Dof* genes in response to stress may show variation.

In *B. napus, BnCDF1* (*BnaDof54* in our study) has been reported to play role in freezing tolerance (Xu and Dai 2016). Based on our expression analysis this gene was highly upregulated upon exposure to 4°C for 24h. Furthermore, the overexpression of two tomato *CDF* genes (*SlCDF1* and *SlCDF3*) in Arabidopsis enhanced drought and salt tolerance (Corrales et al. 2014). The overexpression of Arabidopsis’ *CDF3* also enhances the tolerance of transgenic Arabidopsis plants to drought, cold and osmotic stress (Corrales et al. 2017). The *CDF3* orthologous genes in *B. napus, BnaDof53* and *BnaDof102* significantly upregulated in response cold stress and slightly in response to heat stress. In comparison to control conditions these two genes showed slight reduction in gene expression under drought. *CDFs* thus play potential role in abiotic stress tolerance, in addition with their role in flowering time control (Renau-Morata et al. 2020).

We further observed few *BnaDofs* (−17, −47, −81, −113) associated with functional terms “oxidation-reduction process” and “response to oxidative stress” suggesting a potential role in ROS mediated signalling. In wheat, some *TaDofs* have been suggested to act as dynamic regulators of ROS clearance pathways based on their response to heavy metal stress (Liu et al. 2020). Dof proteins are also involved in phytohormone signalling pathways (Lau and Deng 2010; Waqas et al. 2020). Our cis-acting element analysis of the *BnaDof* promoters and functional annotation of BnaDof proteins revealed the presence of several cis elements responsive to in auxin, abscisic acid, salicylic acid, gibberellic acid, and methyl jasmonate. ABRE elements were enriched in the promoters of 77 *BnaDofs*. ABA dependent pathways and phytohormone signalling are known to be involved in response to abiotic stresses (Agarwal and Jha 2010; Dar et al. 2017; Wani et al. 2016). This suggests that BnaDofs expression gets activated by hormones and they might play role in stress signalling pathways.

The expression analysis exemplifies the stress-responsive nature of *BnaDofs*. The differential regulation of *BnaDofs* may regulate downstream genes involved stress response and probably imparting tolerance. Different *BnaDofs* can be stress-responsive in different tissues and at different developmental stage due to preferential tissue expression of *BnaDofs*. Overall, this study provides a comprehensive understanding of the molecular structure, evolution, and potential functions of *BnaDofs*.

## Conclusions

A systematic analysis of *B. napus* Dof Transcription Factor gene family identified a total of 117 *BnaDofs*. The *BnaDofs* were classified into 9 groups: A, B1, B2, C1, C2.1, C2.2, C3, D1 and D2 based on the phylogenetic analysis. Based on the orthology, synteny and evolutionary analysis and the calculated divergence times indicated that the divergence of *Brassica* and Arabidopsis genus (~17Mya), the whole genome triplication event (9-15Mya) and the formation of *B. napus* (7500 years ago) drove the expansion of *BnaDof* gene family. Synteny analysis also highlighted that majority of the *Dof* genes with known chromosomal locations in *B. napus* did not undergo translocations. The Ka/Ks ratio of the duplicated gene pairs indicated that the *BnaDof* gene pairs underwent purifying selection. Further understanding of the molecular evolutionary mechanism is required to understand how gene duplications, gene loss and rearrangements can lead to expansion of gene families and possible neo-or sub-functionalisation of genes. Tissue-specific expression highlighted the role of *BnaDofs* in organ development and other developmental processes. Most of the *BnaDofs* were responsive to temperature fluctuations and were differentially regulated particularly by the cold stress. Additionally, molecular characterisation, functional annotation and cis-acting element analysis have provided a starting point for further research investigations. Our study support for the involvement of *Dof* gene family in developmental processes and multiple abiotic stress response. Further research is warranted for the dissection of the role of *BnaDofs* and exploring these transcriptional regulators for developing climate change resilient varieties with desirable physiological and agronomic traits.

## Declarations

### Availability of data and materials

All data generated or analysed during this study are included in this article and its supplementary information files Details and accession numbers of the RNA Seq data libraries downloaded from NCBI sequence Read Archive are outlined in Table S9. The expression maps of *BnaDofs* downloaded from Brassica Expression DataBase (BrassicaEDB) are accessible via the following link https://jmp.sh/odBHlfT.

## Authors’ contributions

NL analysed the data and wrote the manuscript with contributions from SB. PLB and MBS conceived the research, supervised, and extensively edited the article.

## Acknowledgements

This research was supported by Melbourne Bioinformatics at the University of Melbourne, project UOM0033. We also acknowledge the support of the ARC Discovery grant DP0988972 and the University of Melbourne Research Scholarship.

