## Supplementary material for "Genome-wide identification and comparative analysis of *Dof* gene family in *Brassica napus*": Figure S1

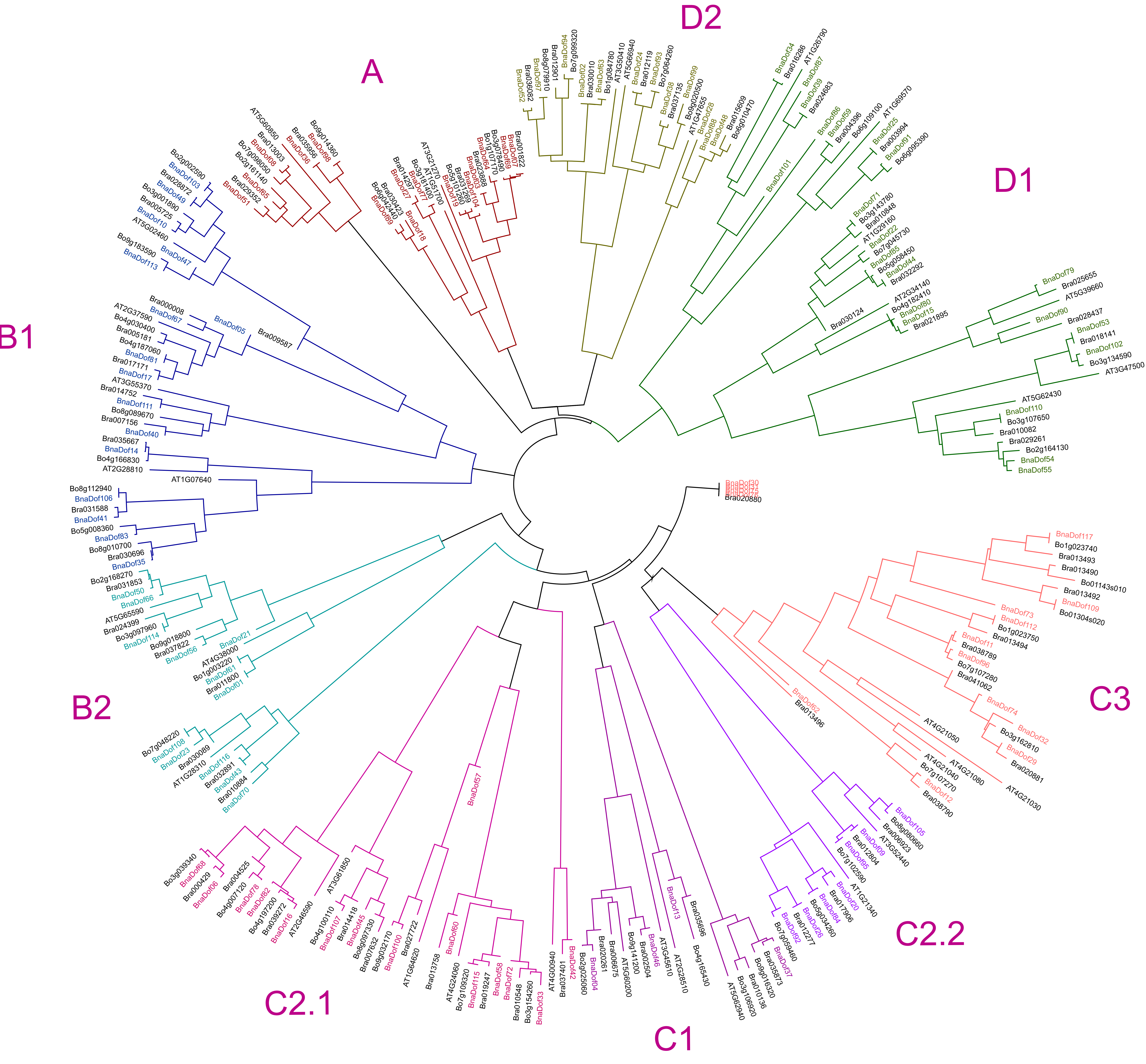


**Figure S1 Phylogenetic tree of Arabidopsis, *B. napus*, *B. oleracea* and *B. rapa* Dof proteins.** The full-length amino acid sequences were aligned using CLUSTALW and the phylogenetic tree using the Neighbor-Joining method was constructed using MEGA7. The percentage of replicate trees in which the associated taxa clustered together in the bootstrap test (1000 replicates) are shown next to the branches. The evolutionary distances were computed using the Poisson correction method. The analysis involved 290 amino acid sequences.
